# Plasmids weaponize conjugation to eliminate non-permissive recipients

**DOI:** 10.64898/2026.02.10.705089

**Authors:** Jonathan N.V. Martinson, Leo C.T. Song, Benjamin E. Rubin

## Abstract

Horizontal gene transfer via conjugative plasmids is a major driver of bacterial evolution. While antibiotic exposure selects for the resistance genes carried by some plasmids, much uncertainty remains in how plasmids persist and spread in microbial communities in the absence of such external selection. Here we show that conjugative plasmids drive their own spread, in part, by using their transfer machinery to enforce a join-or-die ultimatum that selectively eliminates non-permissive recipients through the process of lethal zygosis – the T4SS-mediated elimination of recipients that fail to establish the plasmid. We found that this contact-dependent killing effectively clears a competitive niche for plasmid donors by turning bacterial immune systems into a fatal liability. Specifically, cells resisting plasmid establishment via CRISPR-Cas or Restriction-Modification systems are selectively killed because they fail to acquire the protective exclusion genes encoded on the plasmid, leading to lethal unregulated transfer. We leveraged this insight to resolve an obstacle in bacterial editing by developing “shielded” transposon vectors that co-deliver exclusion genes, increasing the efficiency of gene editing by orders of magnitude. Our results reveal the role of “coercive assimilation” in horizontal gene transfer, exposing a trade-off where the advantage of bacterial immunity to foreign DNA is counterbalanced by lethal zygosis.

## Introduction

Conjugative plasmids drive many aspects of bacterial evolution, yet their persistence in the absence of external selection represents a theoretical conundrum known as the “plasmid paradox” (1). Evolutionary models predict that plasmids should be eliminated from populations via negative selection due to their often high metabolic costs and that beneficial plasmid-encoded genes should migrate to the chromosome, leading to plasmid loss. Although several solutions to this paradox have been proposed (infectious transmission, fluctuating selection, etc.), the precise mechanisms by which plasmids actively invade and persist in uninfected populations remain undercharacterized.

To effectively spread, plasmids must prevent futile self-transmission. When cells carry a conjugative plasmid, they express plasmid-encoded genes that prevent the redundant acquisition of the same plasmid (2,3). This process, known as “plasmid exclusion”, is primarily mediated by entry-exclusion genes (and, to a lesser extent, surface-exclusion genes) which encode membrane proteins that inhibit the formation of productive mating pairs and inhibit DNA transmission (Figure 1A) (3). Exclusion is critical for plasmid fitness, as conjugation is metabolically expensive and redundant delivery is wasteful. However, plasmid exclusion is vital for the host, as well. If exclusion is absent, the Type IV Secretion System (T4SS) continues to engage recipients, leading to multiple simultaneous mating events. This results in death via membrane damage and/or an SOS response triggered by the uncontrolled influx of DNA (Figure 1A) (4). This conjugation-induced killing process, first described by Alfoldi, Jacob, and Wollman in 1957, is known as lethal zygosis (5). While research has focused on the mechanism of lethal zygosis using simplified experimental systems (4–7), its broader implications for horizontal gene transfer and microbial editing remain largely unexplored.

**Figure 1.**
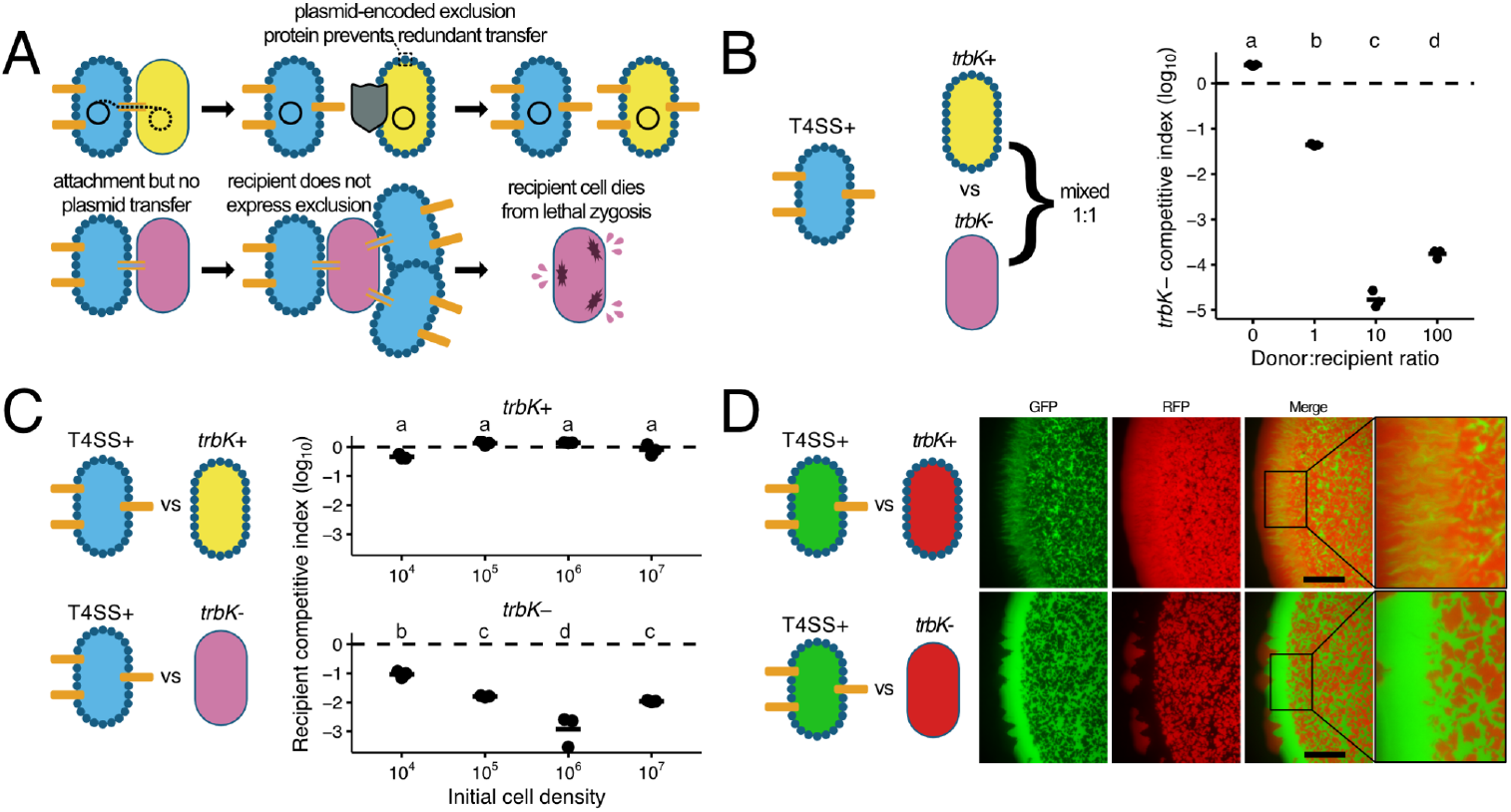
In the absence of plasmid transfer, lethal zygosis eliminates recipients via a density- and ratio-dependent mechanism. **(A)** Diagram of plasmid conjugation and lethal zygosis. **Top:** In permissive recipients (yellow oval), plasmid establishment drives expression of the entry exclusion factor TrbK (dark blue circles), creating a shielded state that blocks subsequent T4SS-mediated donor (light blue oval) contacts. Bottom: In situations where a cell expresses T4SS machinery, but does not transfer plasmid DNA, unshielded recipients are susceptible to repetitive, T4SS punctures (orange rectangles) (lethal zygosis). **(B)** Ratio-dependent killing. Left: Assay schematic. Plasmid-free donors (DATC) expressing chromosomal T4SS genes were mixed with a 1:1 population of shielded (*trbK+*) and unshielded (*trbK-*) recipients. Right: Competitive index of the unshielded population (*trbK*-) relative to the shielded (*trbK*+) population mixed with varying amounts of plasmid-free donors. The dashed line indicates neutrality (CI = 0). Statistical significance was determined by one-way ANOVA with Tukey’s multiple comparisons test. Different letters (a d) indicate statistically significant differences (p < 0.05). Circles represent the competitive index of replicates, bars represent the mean (n = 3). **(C)** Density-dependent elimination. Left: Assay schematic. Plasmid-free donors were mixed 1:1 with either shielded (*trbK*+) or unshielded (*trbK*-) recipients across a range of initial cell densities (10^4^ - 10^7^ cells). Right: Unshielded recipients (*trbK-*, bottom) display reduced fitness (competitive index) at high cell densities compared to shielded recipients (*trbK+*, top). Statistical significance was determined by two-way ANOVA with Tukey’s multiple comparisons test. Different letters (a d) indicate statistically significant differences (p < 0.05). Shielded recipients (*trbK+*) showed no significant differences across densities, but retained significantly higher fitness than *trbK*-recipients at all tested densities (p < 0.05). **(D)** Spatial exclusion. Fluorescence microscopy of competition between GFP-labeled T4SS+ donors (green) and RFP-labeled recipients (red). Shielded recipients (*trbK+*, top) intermix with donors, whereas unshielded recipients (*trbK-*, bottom) are spatially displaced. Scale bar, 750 µm.

In this study, we hypothesized that recipients with DNA defense systems active against conjugative plasmids are susceptible to lethal zygosis, as the degradation of incoming plasmid DNA prevents the expression of protective exclusion proteins. Consistent with this, we found that recipients carrying either a Restriction-Modification (RM) system or CRISPR system targeting the broad host range plasmid RK2 had significantly reduced fitness relative to competing recipients without defense systems. These results suggest that conjugative plasmids function as an expansionist force enforcing a join-or-die ultimatum: recipients must accept the plasmid to gain the protective “shield” of exclusion genes, or resist and face elimination. By selectively killing cells that are non-permissive to plasmid uptake, donors effectively clear a competitive niche – a strategy we term “coercive assimilation.” We applied this insight to microbial editing, reasoning that standard non-replicative delivery vectors likely trigger lethal zygosis because they lack exclusion genes. We engineered a “shielded” delivery system carrying the RK2 entry exclusion gene and demonstrated that this simple addition significantly improves editing efficiency for the randomly integrating Mariner transposon.

## Results

### Lethal zygosis is ratio- and density-dependent

To validate the role of conjugation machinery in driving lethal zygosis, we established a competitive fitness assay comparing recipients with and without the RK2 plasmid entry-exclusion factor, TrbK (8). To isolate the physiological effects of the mating pair formation complex from secondary effects of plasmid transfer on the recipient, we used the strain DATC, which carries conjugation machinery (RK2) integrated into its chromosome (9). Importantly, this strain does not transmit DNA unless transformed with a plasmid carrying an origin of transfer (oriT). This system allowed us to decouple the mechanical process of conjugation from DNA transfer, isolating the specific impact of entry exclusion on recipient fitness.

We first examined the ratio (i.e., frequency) dependence of lethal zygosis by mixing recipients with or without RK2’s entry exclusion gene *trbK* at a 1:1 ratio and competing them in the presence of increasing concentrations of the plasmid-free donor, DATC (Figure 1B). Similar to other studies (6,7), we observed a significant relationship between donor-to-recipient ratio and the fitness of recipients lacking *trbK* (one-way ANOVA, F(3,8) = 1507, p < 0.001). While the competitive index of the *trbK*-population remained close to neutral in the absence of donors (i.e., ∼0), it declined precipitously as the donor-to-recipient ratio increased, reaching a minimum at a 10:1 ratio (Figure 1B). Interestingly, we observed a significant recovery in fitness at the highest donor ratio of 100:1 (Tukey’s HSD, p < 0.001, 100:1 vs 10:1), which may suggest additional physiological stresses at high cell densities. These results confirm that the conjugation machinery alone exerts a potent negative fitness effect on recipients unable to block redundant conjugation events.

Because conjugation is a contact-dependent process, we hypothesized that lethal zygosis would also be governed by cell density, a relationship that, to our knowledge, has not been previously demonstrated. To test this, we performed competition assays between DATC donors and either *trbK*+ or *trbK*-recipients across a range of initial cell densities (10^4^ -10^7^ cells), while holding the initial donor-to-recipient ratio constant (1:1) (Figure 1C). We observed a significant interaction between recipient genotype and cell density (two-way ANOVA, F(3,16) = 32.48, p < 0.0001), confirming that crowding selectively impacts populations without *trbK*. While the presence of the exclusion gene provided protection to *trbK*+ recipients across all densities (Figure 1C, top panel), *trbK*-recipients exhibited significant fitness defects as cell density increased, with peak lethality observed at an initial cell density of 10^6^ (Figure 1C, bottom panel). Similar to the results of the experiments in which donor:recipient ratio was varied (Figure 1B), a recovery in fitness was noted at the highest density (initial density at 10^7^ vs 10^6^ cells; Tukey’s HSD, p < 0.05), potentially reflecting physiological constraints on conjugation in high-density populations.

Finally, we visualized colony-scale spatial dynamics of this antagonism using fluorescence microscopy. We mixed GFP-labeled DATC donors with RFP-labeled recipients on solid agar. When donors were mixed with protected recipients (*trbK+*), the populations intermixed freely. However, mixtures with unprotected recipients (*trbK*-) revealed distinct spatial segregation, characterized by a sharp boundary between the donor and recipient populations (Figure 1D) similar to those observed in other contact-dependent killing weapons (e.g., Type VI secretion systems, CDI) (10,11).

### Bacterial immunity creates a fatal liability during conjugation

DNA defense systems, such as RM and CRISPR-Cas, are often characterized as fitness-enhancing elements that protect cells from hostile mobile genetic elements (MGEs) like bacteriophage (12). However, avoiding lethal zygosis from a conjugative donor requires receipt of the plasmid and expression of the encoded exclusion genes. We reasoned that if intracellular immune systems degrade the incoming DNA before these genes can be expressed, active defense would create a fatal vulnerability: by blocking plasmid establishment, the host fails to acquire the protective “shield” and is consequently eliminated by lethal zygosis (Figure 2A).

**Figure 2.**
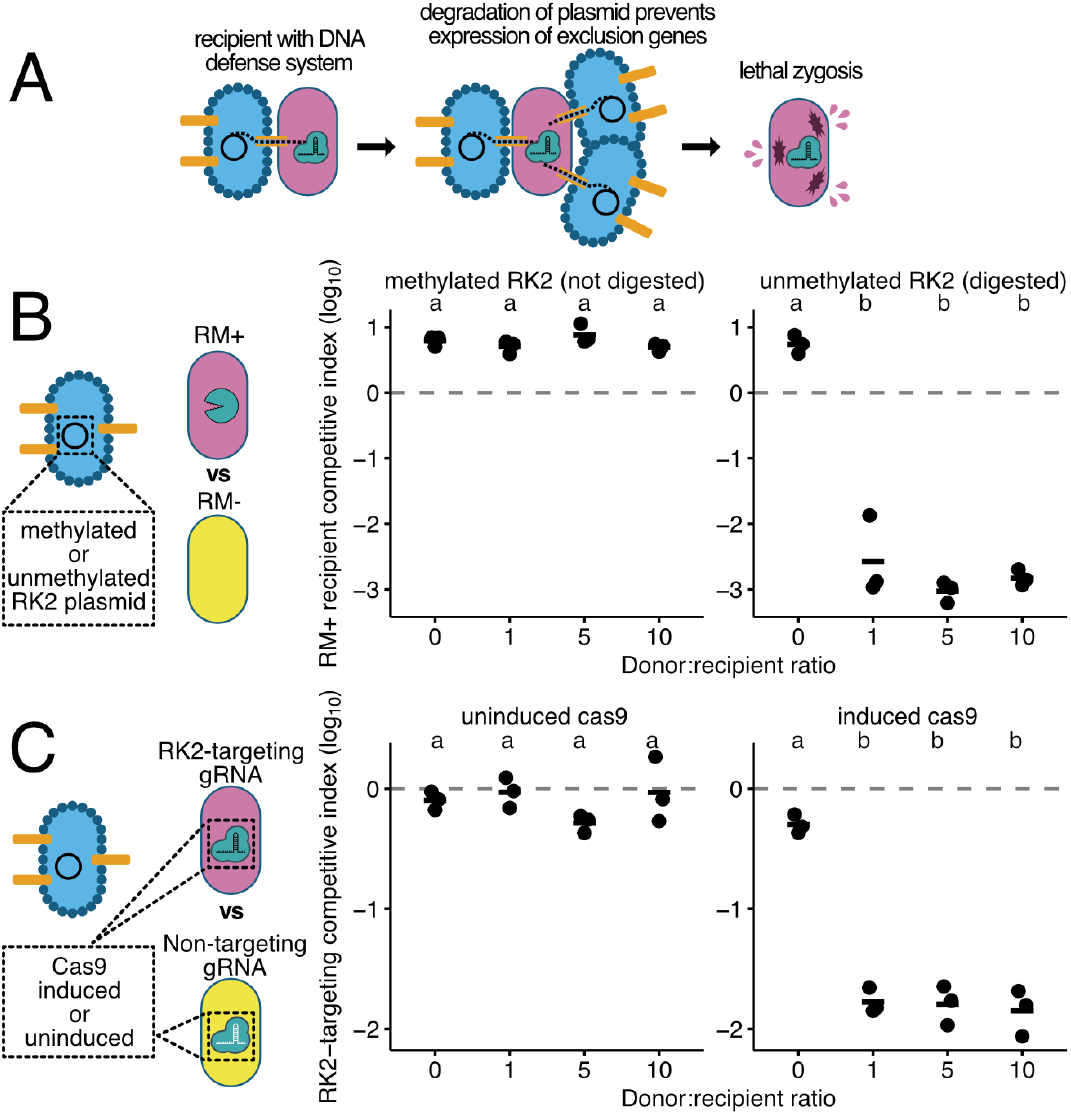
Bacterial immune systems prevent acquisition of exclusion genes, leading to non-permissive recipient elimination via lethal zygosis. **(A)** Model of coercive assimilation. In non-permissive recipients (pink oval), intracellular defense systems (e.g., CRISPR-Cas9, teal) degrade the incoming plasmid, preventing TrbK expression. The unshielded recipient remains susceptible to repetitive, T4SS punctures and uncontrolled DNA delivery (lethal zygosis). **(B)** Restriction-Modification (defense). Left: Schematic of the RM competition assay. RM+ recipients (non-permissive, pink) and RM-recipients (permissive, yellow) compete in the presence of varying amounts of donor cells carrying RK2 plasmids (blue) that are either methylated (protected from digestion) or unmethylated (susceptible to digestion). Right: Competitive index of RM+ recipients relative to RM-recipients in the presence of donors with methylated (left facet) or unmethylated (right facet) RK2. Statistical significance was determined by two-way ANOVA with Tukey’s multiple comparisons test. Different letters (a - b) indicate statistically significant differences within and between facets. When donors carried unmethylated RK2, RM+ recipients suffered significant fitness costs compared to the methylated plasmid treatment (p < 0.05) **(C)** CRISPR-Cas defense. Left: Schematic of the CRISPR competition assay. Recipients with Cas9 and an RK2-targeting gRNA (non-permissive, pink) or a non-targeting gRNA (permissive, yellow) were competed in the presence of varying amounts of RK2 donor. Right: Competitive index of targeting recipients relative to non-targeting recipients under uninduced or induced conditions. Different letters (a - b) indicate statistically significant differences (p < 0.05). Circles represent the competitive index of replicates, bars represent the mean (n = 3).

To test this hypothesis, we first investigated the effects of the RM system, EcoRII, that prevents establishment of RK2 during conjugation (13). We competed RM+ recipients (non-permissive to plasmid replication) against recipients lacking the restriction enzyme (permissive). Crucially, we modulated the “visibility” of the incoming plasmid to the RM system by using donors carrying either methylated (invisible) or unmethylated (visible) RK2 plasmids. We observed a significant interaction between donor methylation status and donor-to-recipient ratio (two-way ANOVA, F(3,16) = 80.07, p < 0.0001), indicating that restriction activity alters the outcome of the competition. When donors carried methylated plasmids, which evade EcoRII digestion, the non-permissive recipients outcompeted the permissive recipients (Figure 2B). In contrast, when donors carried unmethylated plasmids, the RM+ recipients were strongly selected against with a ∼1000-fold reduction in fitness in the presence of donor cells (Tukey’s HSD, p < 0.05 vs methylated donor treatment).

To determine if this effect extended to other defense systems, we performed a competitive fitness assay with recipients carrying an inducible CRISPR-Cas9 (Figure 2C). We competed recipients expressing either a non-targeting guide RNA (gRNA; permissive) or a gRNA targeting the RK2 origin of replication (non-permissive) in the presence of varying amounts of RK2 donor cells. We observed a significant interaction between Cas9 induction and donor-to-recipient ratio (two-way ANOVA, F(3,16) = 37.70, p < 0.0001). Consistent with the RM results, the fitness of the non-permissive recipient population collapsed in the presence of donors when Cas9 was induced with a nearly 100-fold reduction in fitness compared to the uninduced treatments (Tukey’s HSD, p < 0.05).

Notably, for these experiments (Figure 2), the magnitude of fitness reduction in the RK2-targeting recipients remained consistent regardless of donor-to-recipient ratio (e.g., ∼ -3 for the RM+ recipients (Figure 2B) and ∼ -1.75 for the RK2-targeting gRNA recipients (Figure 2C)). These results were distinct from those observed in the plasmid-free donor experiments, where fitness of recipients without exclusion genes changed depending on the donor-to-recipient ratio (Figure 1B) or the initial cell density (Figure 1C). We reasoned this discrepancy could be potentially driven by (1) Toxin-Antitoxin (TA) activation, where a plasmid cleavage event triggers a lethal-zygosis-independent killing by destroying genes encoding short-lived antitoxins; or (2) secondary donor saturation, where the rapid conversion of permissive competitors into new donors artificially inflates the effective donor-to-recipient ratio.

To disentangle these possibilities, we used DATC to deliver a minimal mobilizable vector containing only the essential components for replication, transfer, and exclusion (RK2’s *trfA*/oriV, oriT, *trbK*, respectively), but lacking TA modules such as *parDE (14)* and the conjugative machinery required for secondary transfer (Supplementary Figure 1A). When we challenged induced CRISPR-Cas9 recipients that targeted the RK2 origin of replication with this non-mobilizable, TA-free vector, fitness effects remained consistent (∼ -1.75) across the donor-to-recipient ratios tested (Supplementary Figure 1B; Tukey’s HSD, p > 0.05 for all comparisons between RK2-targeting ratios) suggesting that TA systems and secondary transfer are insufficient explanatory variables. Alternatively, the differences in donor-to-recipient ratio impact observed between the plasmid-free donor trials (Figure 1B) and the immune-competent recipient experiments (Figure 2) may stem from the SOS response triggered during DNA transfer. Unlike the donors in Figure 1B, the RK2 donors in Figure 2 actively deliver DNA to recipients; this redundant DNA delivery, combined with membrane damage, are known inducers of lethal zygosis (4).

Contact-dependent weapons, such as T6SS, often provide bacteria the capacity to invade communities when initially rare (11). To test if coercive assimilation provides a similar advantage, we competed RK2 donor strains (with or without EcoRII methylase) against an EcoRII+ recipient at varying initial ratios (Supplementary Figure 2A). When donors carried the methylated (protected from digestion) plasmid, fitness was neutral across all ratios, resulting in efficient plasmid transmission (50–100% transconjugants; Supplementary Figure 2B-C). In contrast, unmethylated donors failed to invade when rare. At starting ratios of 1:100 or 1:10, donor fitness decreased nearly 10-fold, and no transconjugants were recovered (Supplementary Figure 2B-C). This suggests that the burden of plasmid carriage outweighs the potential fitness advantages of lethal zygosis when the donor is rare. However, this dynamic inverted when the initial ratio was 1:1, as unmethylated plasmid donors’ fitness increased nearly 3 orders of magnitude (Supplementary Figure 2B). These data suggest that the deadly effects of defense system-induced lethal zygosis exhibit positive frequency-dependence.

### Exclusion genes improve genetic editing efficiency

Delivery of non-replicative plasmids (i.e. suicide plasmids) is essential for forward-genetics approaches such as transposon mutagenesis, allelic exchange mutagenesis, and potentially *in vivo* microbiome editing, where the use of replicative vectors encoding editors raises concerns about uncontained dissemination of MGEs (15–17). However, the delivery of non-replicative vectors via conjugation often results in low efficiencies, especially in non-model organisms (17,18). We hypothesized that this poor performance is partially attributable to lethal zygosis. Because suicide plasmids lack exclusion genes, they render transconjugants susceptible to uncontrolled conjugation and subsequent death. To test this, we integrated RK2’s entry exclusion gene, *trbK*, into the payload of the randomly integrating Mariner transposon system. We reasoned that by providing recipient strains a “shield” against lethal zygosis in the form of an exclusion gene, we would increase editing efficiency by preventing conjugation-induced death (Figure 3A).

**Figure 3.**
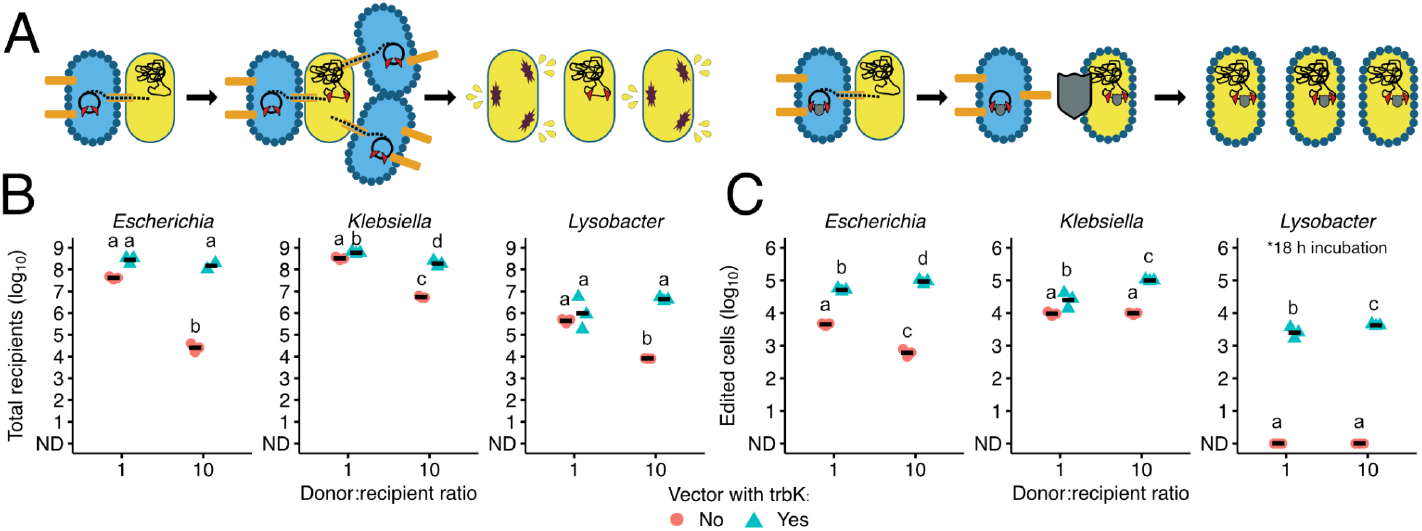
Inclusion of exclusion genes shields recipients from lethal zygosis and improves transposon editing yield. **(A)** (Left) Schematic illustrating the mechanism of lethal zygosis during standard transposon delivery (left) versus the proposed model of how an exclusion gene in a transposon payload delivered improves editing yield by shielding cells. **(B)** Recipient survival (total viable recipients) and **(C)** Editing efficiency (total transposed, antibiotic-resistant mutants) following conjugation with *Escherichia, Klebsiella*, and *Lysobacter* recipients. Different letters denote statistically significant differences (p < 0.05, two-way ANOVA with Tukey’s HSD). Note: All conjugations were performed for 5 hours, with the exception of *Lysobacter* editing assays (C), which required 18 hours to yield detectable mutants (no edits were observed at 5 hours). Symbols: Blue triangles: transposon payload with *trbK*; Red circles: transposon payload without *trbK*. Horizontal bars indicate the mean (n = 3).

We found that delivering standard vectors (lacking *trbK*) at high donor-to-recipient ratios (10:1) caused a significant reduction in the number of viable recipients across *Escherichia, Klebsiella*, and *Lysobacter* populations (Figure 3B). This lethal effect was effectively neutralized when the *trbK* gene was co-delivered in the Mariner transposon payload; the “shielded” vectors maintained high recipient viability at both 1:1 and 10:1 ratios (p < 0.05, two-way ANOVA with Tukey’s HSD). This protective effect translated into improved editing yields, with *trbK-*containing vectors consistently generating significantly more antibiotic-resistant mutants (Figure 3C, p < 0.05, two-way ANOVA with Tukey’s HSD). This advantage was most notable in *Lysobacter*: while a 5-hour conjugation yielded no mutants, extending the assay to 18 hours resulted in edited cells exclusively with the *trbK*-shielded vector. Notably, the “shielding” effect in *E. coli* at the 10:1 ratio resulted in a 38,000-fold improvement in recipient survival while increasing the editing yield by ∼150-fold. Given that the increase in survival was orders of magnitude higher than the increase in editing, we conclude that *trbK*, transiently expressed from the delivery vector without integration, functions as a population-wide shield against lethal zygosis, preserving the recipient pool for subsequent modification.

## Discussion

In this study we demonstrate that plasmids weaponize conjugation to cull recipients that resist plasmid acquisition. Specifically, we show that when cells mount an active defense against plasmids, they are sensitized to lethal zygosis because they degrade the DNA encoding exclusion proteins that protect cells from uncontrolled, repeated conjugation events (Figure 2). In this way, conjugative plasmids provide a strong selective pressure for their own engraftment.

This mechanism creates a tradeoff whereby bacterial immune systems that are often critical for surviving lytic phage pressure (12) become a liability during conjugation. These countervailing forces could feasibly establish a cyclic rock-paper-scissors dynamic (19): Phage pressure selects for immunity; immunity sensitizes bacteria to lethal zygosis, increasing population permissiveness; and this permissiveness resets the cycle by exposing the population to phage predation once again (Figure 4). Although this is a simplified model, it provides a mechanistic rationale for the observation that CRISPR systems target plasmids far less frequently than bacteriophage, despite the metabolic burden plasmids impose (20). If targeting a plasmid induces lethal zygosis, anti-plasmid spacers would be actively selected against. Consistent with this, restriction–modification systems can restrict phage with orders of magnitude more efficacy than conjugative plasmids (13,21). While these observations do not imply causation, they align with a model in which coercive assimilation creates a selective pressure against plasmid targeting while preserving phage defense.

**Figure 4.**
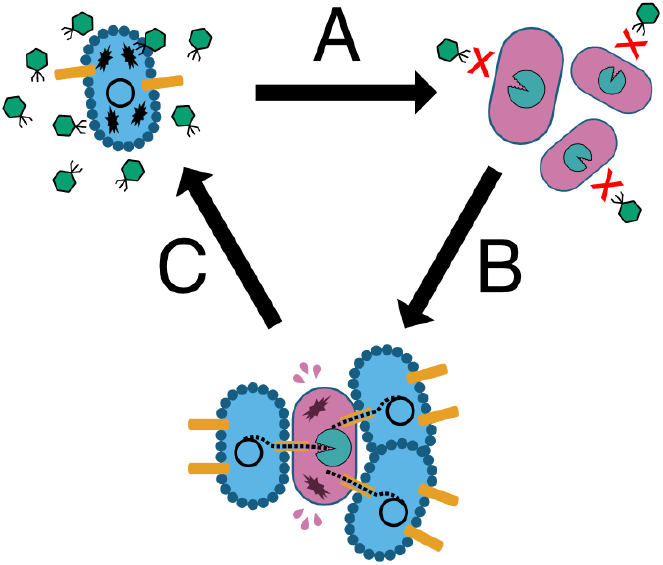
A “Rock-Paper-Scissors” evolutionary dynamic driven by the incompatibility of defense and conjugation. **(A)** Phage pressure selects for immunity by eliminating susceptible permissive hosts. **(B)** Lethal zygosis selects against immunity, as cells lacking active defense survive plasmid uptake and acquire exclusion genes. **(C)** This permissive state resets the cycle by exposing the population to phage predation once again.

Coercive assimilation’s relevance to plasmid biology likely extends beyond RK2 and *E. coli*. Historically, the study of plasmid exclusion and lethal zygosis has focused on the F plasmid (2,5,6,22), though exclusion genes are ubiquitous across conjugative systems (3). For example, lethal zygosis has been observed in *Bacillus subtilis* following the loss of the ICEBs1 exclusion gene, *yddJ* (23,24). Additionally, recent work has shown that RK2 conjugation machinery is capable of causing interspecies lethal zygosis against diverse members of Enterobacteriales and Pseudomonadales (7). Thus, coercive assimilation may serve as a generalized strategy employed by MGEs across the bacterial tree of life.

We propose that the use of lethal zygosis to kill non-permissive competitors is not accidental, but rather a density-dependent competitive strategy. Although lethal zygosis is ineffective as an offensive weapon when donors are rare (Supplementary Figure 2), it may serve as a potent means of colonization resistance when a donor population is dominant, allowing established populations to actively prevent encroachment of non-permissive competitors in their niche. This is particularly relevant in high-density ecosystems such as biofilms, where frequent cell-to-cell contact facilitates conjugation (and presumably lethal zygosis). Furthermore, this mechanism likely functions in concert with other plasmid-encoded arsenals like diffusible toxins (e.g., bacteriocins). Recent experimental and modeling work has found that the combination of contact-dependent and contact-independent weapons significantly enhances a strain’s competitive ability to defend and invade a niche (11). Together, these data suggest that lethal zygosis may shed light on the mechanisms of plasmid maintenance in the absence of external selection (i.e. the “plasmid paradox”) by providing a selective advantage to plasmid carriers under high-density conditions.

Finally, we applied our findings to synthetic biology. Standard suicide vectors inadvertently mimic the conditions for lethal zygosis: they engage the T4SS to deliver DNA but fail to establish the plasmid, thereby condemning recipients to death. To resolve this, we sought to improve conjugal delivery by engineering a Mariner transposon to co-deliver the RK2 exclusion gene, *trbK*. Providing this “shield” not only improved transposition efficiency but also protected recipient populations from lethal zygosis (Figure 3). This strategy proved critical for the soil bacterium *Lysobacter* sp., where standard vectors failed to yield a single mutant but successful editing was achieved when the recipient was protected by *trbK*. These results suggest that some microbes deemed “intractable” may not be recalcitrant to genetic tools, but rather hypersensitive to lethal zygosis induced by their delivery. The incorporation of exclusion genes into suicide editing vectors could improve our ability to edit previously intractable species.

This study has several limitations that warrant further investigation. Our work focused on the broad host range RK2 plasmid with hosts within the Gammaproteobacteria. Plasmids with narrower host ranges, stricter conjugation regulation, or anti-defense systems (e.g., anti-CRISPRs, anti-restriction proteins) may exhibit distinct ecological dynamics (25), and the extent of lethal zygosis could differ substantially. Additionally, we performed our competition assays under simplified, nutrient-rich, aerobic conditions that do not fully recapitulate the metabolic and ecological stresses cells face in natural environments. Finally, we examined only immune systems that recognize and target specific DNA motifs (CRISPR-Cas and RM).

Defense systems that respond to other signals like changes to the cell membrane or intracellular metabolite pools may impose different selective pressures (26,27). More broadly, immune system liabilities are not unique to conjugation; defense systems targeting temperate phage can induce lethal autoimmunity, significantly reducing lysogen fitness, and in some circumstances, select for immune system loss (28). Future work should probe coercive assimilation across a broader diversity of organisms and within their native microbiomes and environments.

Plasmids have long been recognized as key vectors of virulence and antibiotic resistance, but our understanding of their ecological impact has been mostly limited to the accessory genes they carry and the metabolic load they impose on hosts (29,30). Our findings expand this view by revealing that conjugative plasmids can act as direct ecological agents, restructuring microbial populations through contact-dependent antagonism. More broadly, understanding how plasmids drive this coercive assimilation to shape the tradeoffs between immunity, permissiveness, and horizontal gene transfer will be essential for predicting microbiome dynamics and developing more effective genetic tools.

## Methods

### Culture conditions

Liquid bacterial cultures were grown with shaking (200 rpm) in Terrific Broth (TB; Fisher Scientific, Cat. No. BP2468500) or lysogeny broth (LB; RPI, Cat. No. L24060-5000.0). For solid media, cells were cultured on lysogeny broth supplemented with 1.5% agar (BD, Cat. No. 214010). Experiments were performed at 37º C or 30º C, as needed. Where appropriate, antibiotics were added to liquid and solid media as follows: carbenicillin (100 µg/mL), chloramphenicol (34 µg/mL), gentamicin (10 µg/mL), kanamycin (25 µg/mL), and spectinomycin (50 µg/mL). Diaminopimelic acid (DAP) was added to media at 3 mM to support the growth of DAP-auxotrophic strains.

### Bacterial strains

The bacterial strains used in this study are listed in Supplementary Table 1. Cloning was primarily performed in *Escherichia coli* EC100D *pir*+ (Epicenter, Cat. No. 75927-934) and to a lesser extent BL21(DE3) (New England Biolabs, Cat. No. C2527H) as it is Dcm-negative (EcoRII modifies the same motif). Delivery of R6K suicide plasmids was performed using the DAP-auxotrophic DATC, which carries the RK2 conjugation machinery and R6K *pir* gene integrated into its chromosome (9). For transposon delivery experiments, the following recipient strains were used: *E. coli* BW25113, *K. michiganensis* M5a1, and *Lysobacter* sp. OAE881.

### Plasmid construction

The plasmids used and generated in this study are listed in Supplementary Table 2. Routine cloning was performed using Gibson assembly (NEBuilder® HiFi DNA Assembly Master Mix, New England Biolabs) from PCR-amplified fragments (Q5® High-Fidelity DNA Polymerase, New England Biolabs). Plasmid assemblies were confirmed by Oxford Nanopore plasmid sequencing (Plasmidsaurus). For all *trbK* containing constructs, the gene was amplified from the RK2 plasmid and expression was driven by the medium-strength constitutive Anderson promoter (BBa_J23108) coupled with a ribosome binding site designed using the “Control Translation” function on denovodna.com, targeting a translation efficiency rate of 50,000 a.u. (31). Plasmid pN3’s EcoRII RM system (NCBI reference sequence NC_015599.1) was synthesized (Twist Bioscience) based on coordinates described previously (32). CRISPR guide RNAs were designed to target RK2’s oriV using the Benchling guide design tool (33). The “shielded” transposon vector, pJNVM0283, was constructed by inserting RK2’s *trbK* inside the transposon repeats of a previously developed Mariner transposon vector (34).

### General Competition and Mating Assays

Unless otherwise noted, all competition and mating assays followed a standardized workflow.

1. Preparation: Cells were grown in TB with appropriate selection. For experiments in Figure 1, stationary phase cells from overnight cultures were used directly. For experiments in Figures 2 and 3, overnight cultures were back-diluted and grown to mid-log phase prior to harvest.
2. Washing & Normalization: Cells were washed three times in phosphate-buffered saline (PBS) at 3500 x g. Recipients were normalized to OD_600_ = 1.0 (see exceptions below). In cases where 2 recipient strains were competed, strains were mixed at a 1:1 ratio. We resuspended cells by pipetting instead of vortexing to avoid damaging the conjugative apparatus. 3. Mating: The recipient mixture was combined with donor strains at varying donor-to-recipient ratios. 10µL of mixtures were spotted onto LB agar (supplemented with DAP for auxotrophs) and dried by a flame. 4. Harvest: After incubation at 37°C for 5 hours, spots were scraped into 100µL PBS using sterile loops, serially diluted, and plated on selective media.

#### Specific experimental variations are detailed below

Ratio-Dependent Lethal Zygosis (Figs. 1B, 2B): Donors were adjusted to varying densities prior to mixing to test ratio effects. In Figure 1B, donor inputs were OD_600_ 1, 10, and 100. In Figures 2B and 2C, donor inputs were OD_600_ 1, 5, and 10.

Density-Dependent Survival (Fig. 1C): Donors and recipients were normalized to OD_600_ 100, mixed 1:1, and serially diluted in PBS prior to spotting to achieve initial spot densities of 10^7^ to 10^4^ cells (total # of cells per spot).

CRISPR-Cas9 Induction (Fig. 2C, Supp Fig 1): Recipient cultures were split into induced (100 ng/µL anhydrotetracycline [aTc]) and uninduced conditions and incubated for 1 hour at 37°C prior to washing. Mating spots were plated on LB agar supplemented with aTc where appropriate.

Donor Fitness from Rare (Supp Fig 2): The donors were the focal strains and, as such, were normalized to OD_600_ = 1 while the recipients were adjusted to OD_600_ 1, 5, and 10 to assess donor:recipient ratios 1:1, 1:5, and 1:10, respectively. Transconjugants were quantified by using multiple antibiotics in LB agar (kanamycin and gentamicin).

Shielded Transposon Mutagenesis (Fig. 3): Donors (OD_600_ 1 or 10) and recipients (OD_600_ 1.0) were mixed at 1:1 or 10:1 ratios. Conjugation spots were incubated at 30°C for 5 (*E. coli* BW25113 and *K. michiganensis* M5a1) or 18 hours (*Lysobacter* sp. OAE881). Cells were plated on selective media (LB+Chloramphenicol) to enumerate transposition and plain LB for total survival.

### Calculation of Competitive Index

To quantify the relative fitness of a focal strain versus a comparison strain, the competitive index was calculated by determining the ratio of the focal strain and comparison strain at the end of the experiment and dividing it by the ratio at the beginning of the experiment. Following this calculation, the results were log_10_ transformed.

### Fluorescence Microscopy

For spatial analysis (Figure 1D), GFP-expressing T4SS+ cells (sJNVM0375) were mixed 1:1 with RFP-expressing strains with or without *trbK* (sJNVM0378 or sJNVM0390) at a low initial density (OD_600_ = 0.1). Mixtures (10µL, approximately 10^4^ cells) were spotted onto LB agar and incubated for 22 hours at 37°C to allow growth. Colonies were imaged using a 4X objective lens on an EVOS M5000 imaging system (Thermo Fisher) using GFP and RFP fluorescence channels.

### Data visualization and statistical analysis

Data visualization and calculations were performed using R v4.4.0 using the tidyverse (35). Competitive index and cell population counts were log_10_ transformed. Statistical comparisons were performed using one-way or two-way ANOVA followed by Tukey’s Honest Significant Difference (HSD) post-hoc test for multiple comparisons. A p-value of <0.05 was considered statistically significant.

## Data availability

Plasmid sequences, maps, code, and raw data will be made publicly available upon peer-reviewed publication. During the review process, materials are available from the corresponding author upon reasonable request.

## Acknowledgements

We express our gratitude to Brady Cress for generously providing us with several plasmids and valuable experimental suggestions. We also acknowledge the support from the Berkeley Initiative for Optimized Microbiome Editing (BIOME), particularly from Jennifer Doudna, Jill Banfield, Brad Ringeisen, Audrey Glynn, and Rachel K. Evans. We also thank Ryuichi Ono, Jaymin Patel, Lindsey Pieper-Zapien, Zachary LaTurner, Susan Abrahamson, Liana Yong, Michael Dills, Bingliang Xie, Aria Kim, Sarah Hasham, and Madeline Hayes for providing experimental suggestions, reagent preparation, and/or text edits. The authors utilized Gemini (Google) for coding assistance and editing the text. However, all code, analyses, and manuscript content underwent checks and verifications by the authors, who bear full responsibility for the work.

We’d like to thank our funders: The Audacious Project: This work was supported in part by Lyda Hill Philanthropies, Acton Family Giving, the Valhalla Foundation, Hastings/Quillin Fund - an advised fund of the Silicon Valley Community Foundation, the CH Foundation, Laura and Gary Lauder and Family, the Sea Grape Foundation, the Emerson Collective, Mike Schroepfer and Erin Hoffman Family Fund - an advised fund of Silicon Valley Community Foundation, the Anne Wojcicki Foundation through The Audacious Project at the Innovative Genomics Institute; Leona M. and Harry B. Helmsley Charitable Trust: This work was funded in part by grant [G-2302-06692] from The Leona M. and Harry B. Helmsley Charitable Trust; Shurl and Kay Curci Foundation: This work was also supported by a Research Award from the Shurl and Kay Curci Foundation (https://curcifoundation.org) to the Innovative Genomics Institute Genomic Tool Discovery Program at UC Berkeley, awarded to B.E.R. Innovative Genomics Institute: This work was supported in part by the Innovative Genomics Institute.

## Competing interests

The Regents of the University of California have patents pending related to this work on which J.N.V.M. and B.E.R. are inventors.

---

**Supplementary Figure 1.**
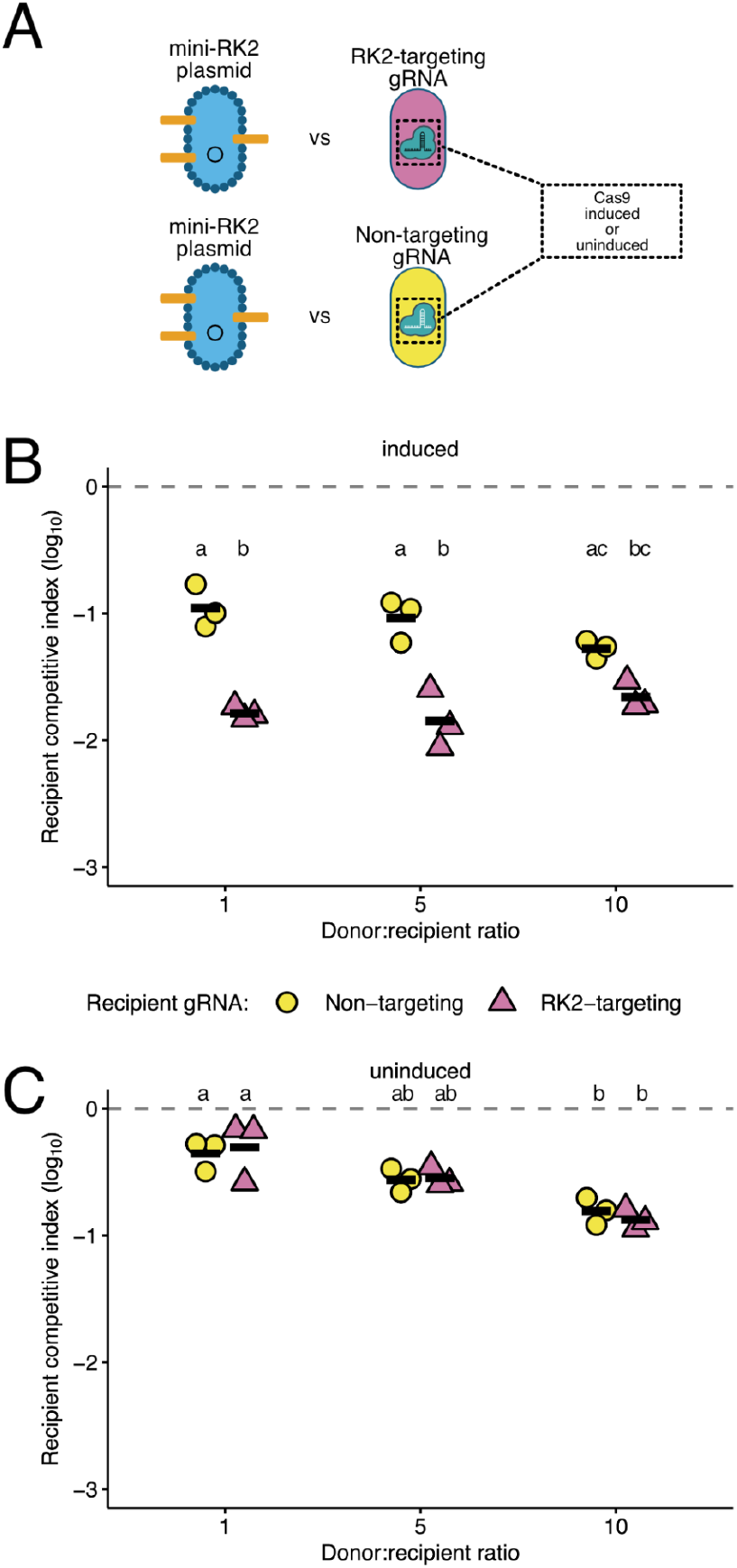
Coercive assimilation is not dependent on TA systems or secondary transfer. **(A)** Schematic of the experimental design. Recipient cells carrying either an RK2-targeting gRNA (pink triangles) or a non-targeting gRNA (yellow circles) were competed against donors carrying a mini-RK2 plasmid that was not self-transmissible and lacked TA systems. **(B, C)** Recipient competitive index across varying donor:recipient ratios (1:1, 5:1, and 10:1) under **(B)** induced or **(C)** uninduced conditions. Individual data points represent biological replicates (n = 3), and black horizontal bars indicate the mean. Different letters denote statistically significant differences (p < 0.05, two-way ANOVA with Tukey’s HSD). In **(B)**, the difference between targeting and non-targeting groups at ratio 10:1 was marginal (p = 0.07), resulting in the shared letter ‘c’. In **(C)**, no significant difference was detected between targeting and non-targeting recipients at any ratio (p > 0.05); the shift from ‘a’ to ‘b’ across ratios indicates a significant main effect of donor density on recipient fitness independent of gRNA identity.

**Supplementary Figure 2.**
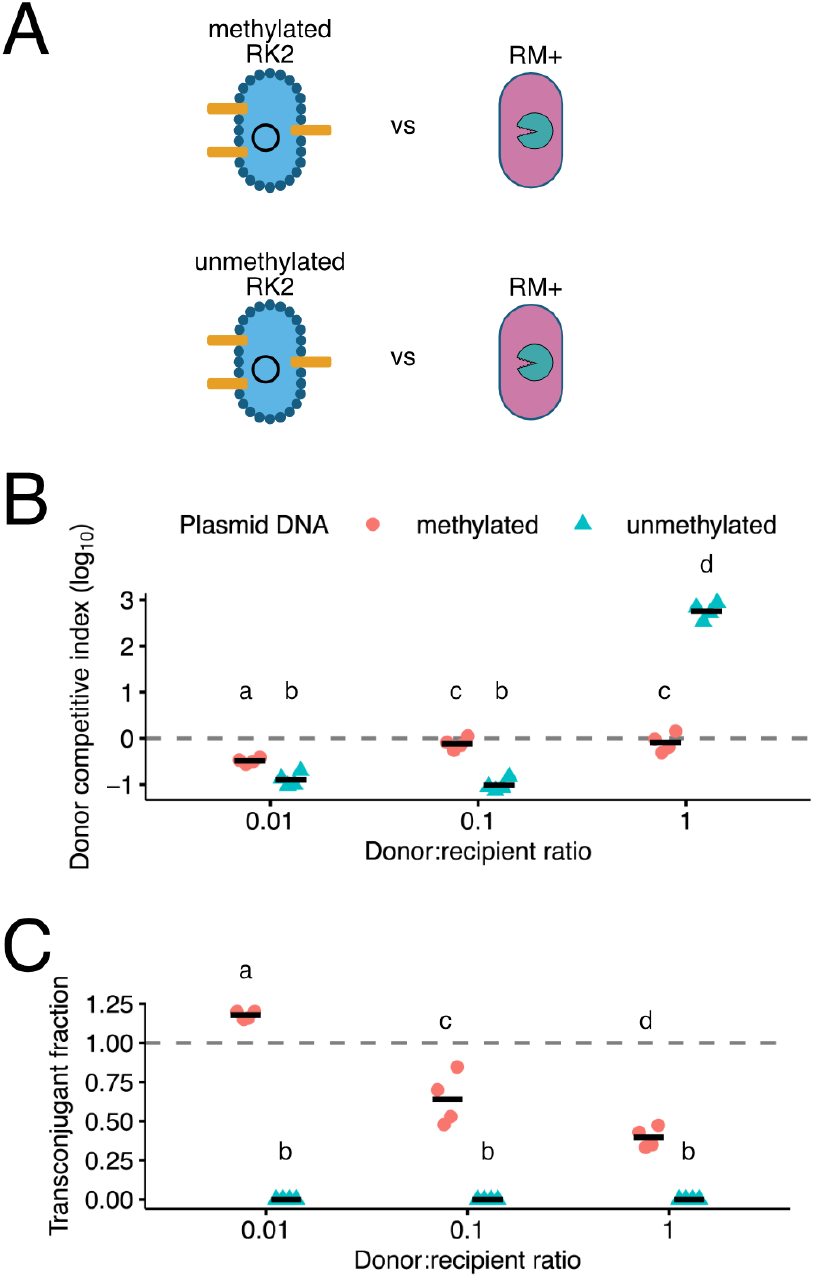
RK2 donor fitness changes depending on donor:recipient ratio and permissiveness of recipient. **(A)** Diagram of experimental setup. RK2 donor cells with (top) or without (bottom) methylase were mixed with recipients with a restriction-modification system (EcoRII) at variable ratios. **(B)** Competitive index of donor population with (red circle) and without (blue triangle) methylation mixed at 1:100, 1:10, and 1:1 donor-to-recipient ratios (n = 4). Different letters denote statistically significant differences (p < 0.05, two-way ANOVA with Tukey’s HSD). **(C)** Fraction of transconjugant recipients when plasmid was methylated or unmethylated.

